# Improving vaccine efficacy based on non-covalent photoinactivation of microorganisms - sources vaccine antigens (MSVA)

**DOI:** 10.1101/2020.07.14.201970

**Authors:** Artur Martynov, Boris Farber, Tatyana Osolodchenko, Ilya Klein

## Abstract

One of the most promising methods for non-covalent inactivation of vaccine-producing microorganisms is the use of photoinactivation using riboflavin derivatives. The study used a dynamic combinatorial derivative of riboflavin - succinyl-maleinyl riboflavin. Corpuscular vaccines are divided into the following groups: from 2AB to 5AB - bacteria were inactivated by riboflavin derivative and blue light, and groups from 6AB to 9AB were inactivated by formalin (0.1% formalin in 9 log CFU was kept for 2 weeks in an thermostat and then sterility was determined - bacterial growth was not observed). A dynamic derivative of riboflavin at a final concentration of 0.02% can photo inactivate 6 time more bacteria P. Aeruginosa and E. coli than riboflavin. The minimum effective blue light emitter power (450 nm) for the photodynamic inactivation of both P. aeruginosa and E. coli is 1024.2 mW / cm2. In groups of mice pre-vaccinated intraperitoneally with corpuscular photo inactivated vaccines based on suspensions of and E. coli at doses of 0.5-1.0 ml 4 log (CFU) / mL, 100% survival of all animals was observed, whereas in control group with the same type of vaccines but formalin-treated vaccines, it failed to achieve a 100% protective effect.

## Introduction

Modern vaccines are one of the most effective approaches to reduce mortality and to control infectious disease in the human population [1][2]. However, a number of problems remain unresolved in the field of vaccine use (efficacy).

When developing vaccines an essential part of the research is not only to confirm their high immunogenicity and the presence of protective properties against infection, the pathogens, but also is necessary to determine the absence of reactogenicity and allergenicity. Both reactogenicity and allergenicity are associated with the presence of allergenic antigens and haptens in the vaccine [3].

These abnormal antigens are formed during the chemical inactivation of the microorganisms which produce vaccine antigens by inactivators such as formalin and propiolactone, as well as mercury preparations such as merthiolate. As a result of a chemical reaction between formalin and amino groups of proteins, Schiff bases are formed, which are then oxidized to formylated amino acid residues [4,5]. At the same time, the antigenicity of the initial proteins changes significantly, and the vaccinated organism synthesizes antibodies not to the original antigen, but to a formylated one [6].

In fact, usually the initial immunogenicity of microbial proteins under the action of formalin decreases twofold or more. Moreover, abnormal formylated proteins, lipoproteins, often act as allergens [7]. Reducing the allergenicity and reactogenicity of antigens is possible, if microorganism-producent vaccine antigens is inactivated by methods that exclude the formation of covalent bonds between proteins and inactivators.

Accordingly, inactivation of MSVA occurs without the formation of new abnormal antigens / allergens. One of the directions in the development of non-reactogenic and non-allergenic vaccines is the replacement of formalin and propiolactone with non-covalent inactivators. This is one of the most promising methods for non-covalent inactivation of MSVA, because is the use of photodynamic inactivation using riboflavin derivatives [8].

Past research has established that riboflavin derivatives have the ability to selectively accumulate in the region of flavin aptamers of DNA / RNA in the cell, without violating the vital functions of the cell and the functioning of proteins [9,10].

In fact, after above method approach, an MSVA inactivated remains with intact proteins, but with destroyed and lysed DNA and RNA [11]. After irradiation of riboflavin a harmless lumiflavin is formed. Lumiflavin is also a metabolite of human tissues and does not require cleaning the vaccine from it (as it used for formalin and merthiolate) [12].

The immunogenicity of proteins - vaccine antigens does not decrease, and allergenicity and reactogenicity does not increase, because the structure of the proteins as a whole remains unchanged. The method of riboflavin inactivation of infectious agents widely used in hematology in blood products for transfusion. This method used for disinfection donated blood on Mirasol devices [12].

It was proven the safety and high inactivating activity against microorganisms of the photoinactivation system. Unfortunately, there are very few publications in regards of the effect of riboflavin inactivation on immunogenicity, allergenicity, reactogenicity and protective properties of vaccines [13]. One of the disadvantages of riboflavin inactivation is low solubility of riboflavin and its low permeability through the microbial membrane. For a number of riboflavin derivatives, an increase in antimicrobial activity under the same conditions by 6 log CFU was previously shown [14].

Based on TRIZ (the Theory of Inventive Problem Solving) and the experience of our predecessors, our company came up with a solution to the problem outlined above, by synthesizing a dynamic self-organizing riboflavin derivative, the activity of which was higher with respect to the conversion of toxin to toxoid. We applied TRIZ-a new philosophy of thinking which is a problem-solving, analysis and forecasting tool derived from the study of patterns of invention evolution and getting generalizable patterns in the nature of inventive solutions. It was discovered several trends (so called Laws of Systems Evolution) that help researchers predict the most likely developments that can be made to a given product. TRIZ has been successfully applied in in huge varieties of different fields all over the world, however, for the design of drugs of new generations at all stages of development. TRIZ could significantly decrease expenses, but has not been used systematically. The application of the principles of TRIZ in pharmaceutical industry and pharmacology opens up broad prospects in the creation of new classes of drugs. Therefore, pharmaceutical industry and pharmacology are huge areas to explore by TRIZ [15].

We utilized Urease, adynamic riboflavin derivative (RD) that has greater solubility (2% or more) and is more active in relation to urease photoactivation upon irradiation of the solution in the visible spectrum at 450 nm within 3 minutes. We have done extensive pioneering work in this area established the safety and high inactivating activity against microorganisms of the photoinactivation system.

Our company came up with solution by synthesizing a dynamic self-organizing riboflavin derivative, the activity of which was an order of magnitude higher with respect to the conversion of toxin to toxoid, using urease as an example [16]. This dynamic riboflavin derivative (RD) had greater solubility (2% or more) and was more active in relation to urease photoactivation upon irradiation of the solution in the visible spectrum at 450 nm within 3 minutes.

The aim of this study was to determine of dependence of the intensity of exposure time by visible radiation on the bacterial and the number of bacteria. This dependence is important for the development of a universal method of inactivation in the development of low-allergenic and non-reactogenic vaccines. Also, this research project includes study of the protective properties of the Pseudomonas corpuscular vaccine, which obtained through photodynamic inactivation using RD (riboflavin derivative) and visible blue light exposure at 450 nm (BL).

We have chosen consciously the use of the visible region of the spectrum. Riboflavin and its derivatives are characterized by two peaks of light absorption - ultraviolet in the region of 250 nm and in the visible region (blue light) at 450 nm [17]. Light of the visible spectrum is not absorbed by water, unlike ultraviolet light [18]. Light of the visible spectrum penetrates not only deep into the nutrient medium, but also through the cell membranes of bacteria [19]. In fact, for light with a wavelength of 450 nm, there are no obstacles to the selective riboflavin photo nuclease activity inside flavin aptamers which may last up to 2-3 minutes.

Existing and known methods of photoinactivation of bacteria require a longer period of processing the solution with ultraviolet light with obligatory mixing of the solution (at least 20 minutes) and more intense radiation of ultraviolet lamps [20]. This requirement due to the low permeability of aqueous solutions for ultraviolet radiation with wavelengths of 220-300 nm.

It was not possible to completely suppress the growth of bacteria, and an increase light exposure is undesirable due to heating effect of the solution. Same time an increase in the concentration of riboflavin (R) is impossible due to its low solubility in water. Greater efficiency of inactivation was observed with a longer exposure (up to 40 minutes) or with a higher concentration of riboflavin and only when ultraviolet radiation was used [21].

The use of ultraviolet radiation is undesirable, especially for a long time, due to the risk of influencing antigenicity and immunogenicity. It happens due to the decarboxylation of external residues of protein amino acids as a result of direct exposure to ultraviolet radiation [20]. Unlike ultraviolet light, visible light penetrates deep into bacterial cells and the turbidity of the solution does not interfere with it. However, our results show quite a strong response and would be a very strong step forward in improving vaccine efficacy.

The aim of our studies was to determine the photoinactivation efficiency of RD of the most common pathogenic bacteria: *P. aeruginosa* and *E. coli* at 450 nm. Goal was also to determine different concentrations of bacteria in solution, and also to confirm the presence of protective properties in the resulting antigens with respect to the development of P. *aeruginosa* and *E. coli* sepsis respectively in mice models. As control model, the same corpuscular antigens were used, but inactivated with formalin according to the standard inactivation protocols.

## Materials and Methods

The methodology of our research was based on experiments with riboflavin and ultraviolet irradiation of bacterial suspensions described in [20].

To test our novel process, we utilized an animal vaccine method of testing. In the experiment, white outbred mice were used, which were kept on a vivarium diet under standard conditions.

The animals were cared for according to the international guidelines GOST 33647-2015 (Principles of the Good Laboratory Practice, GLP), the international recommendations of the European Convention for the Protection of Vertebrate Animals used for Experimental and Other Scientific Purposes (The European Convention, 1986). The protocol of the experimental study was approved by the Ethics Committee of Mechnikov Institute of Microbiology and Immunology of National Academy of Medical Sciences (protocol № 3-2018 of 14.12.2018).

Animals were divided into 18 groups of 20 animals per group. The first two control groups 1A and 1B were injected intraperitoneally with suspensions of live bacteria at a dose of 0.5 ml (9 log CFU) / mL of P. aeruginosa and E. coli, respectively. The remaining 16th groups of animals were administered the appropriate amount of vaccine (inactivated bacteria) preliminarily, 2 weeks prior infection with bacteria.

Inactivated RD-BL vaccines were administered to animals of groups 2AB - 5 AB (see Table 1), and animals of groups 6AB - 9AB were given vaccines inactivated with formalin (0.1% formalin in 9 log CFU was kept for 2 weeks in the thermostat and sterility was determined - bacterial growth was not observed).

**Table 1.** Survival fraction of *E. coli* and *P. aeruginosa* with different BL energy in the BL- RD group (irradiation time: 3 min)

| Power density (mW/cm<sup>2</sup>) | Survival fraction of *E. coli* (%) | Survival fraction of *P. aeruginosa* (%) (B) |
| --- | --- | --- |
| 0 | 100 | 100 |
| 912,5 | 12,3±2,9 | 22,5±2,4 |
| 1024,2 | 1,3±0,4 | 3,5±0,5 |
| 1111,8 | 0 | 0 |
| 1209,9 | 0 | 0 |

## Materials

### Bacteria Isolation and Culture

XMR-strain *Pseudomonas aeruginosa* XMR IMI 047 and isolate XMR-strain of *Escherichia coli XMR* IMI 218 were selected for study. All bacterial strains isolated from patients with post-surgical complications were provided by the Department of Surgery of Kharkov National Medical University, Kharkov State Hospital N 17. The bacterial strains stored in the glycerol tube, then were inoculated in a blood culture dish and resuscitated two times to a logarithmic growth phase. Then the bacterial colonies were scraped with an inoculation loop and placed into the bacterial diluents. After sufficient mixing, 0.5 McFarland (MCF) turbidity standard bacterial suspensions were prepared using a turbid-meter (approximate bacteria concentration of 1∼2 x 10^9^ colony forming units/mL, CFU/mL), which was used for the subsequent experiments.

### Blue Light Source and Photosensitizer

The blue light (BL) irradiation was performed with the LED point source (Luxeor-LC, Dealdent Co., Ltd., Kharkiv, Ukraine) with a wavelength of 450 nm. Calibration of the light energy was carried out before each experiment to ensure that the output power density was in range of 900 mW/cm^2^∼1200 mW/cm^2^. The light power density was measured by optical power meter (Optical Power Meter ST805C PPM FTTX PON, Shandong Senter Electronic Co., Ltd., Shandong, China). The diameter of light spot was about 7 mm. In this experiment, the PS was riboflavin (Sigma-Aldrich Technology Co., Ltd., USA) and synthesized derivative (IV) or (RD) as shown in [15], that were dissolved into 0,02% wt solution with sterile phosphate buffered saline (PBS) and stored at 4°C in dark area.

## Methods

### Qualitative Observation of the Inhibition Effect on Bacteria

*In Vitro*. Stock solution 0,2% wt of riboflavin, and RD (Riboflavin dynamic derivative – RD), was diluted to 0.1% wt with sterile PBS. In the experiment, four different samples were prepared which BL-RD treated sample (Group 1), control treated with BL alone (Group 2), control treated with RD alone (Group 3), and control untreated (Group 3). 200 *µ*L bacterial suspension (0.5MCF) was aseptically transferred into the 9 cm blood agar plates with properly labeled test positions of control samples and BL-RD treated sample. 10 *µ*L diluent RD solution (0.1% wt) was added in each labeled test position of BL-RD treated sample and RD alone treated control sample. Before exposure to BL, the samples to be irradiated were incubated in dark for 20 min. 450 nm BL was used to irradiate BL-RD treated sample and BL alone treated sample. The samples were irradiated by different power of BL light (912,5 mW/cm^2^, 1024,2 mW/ cm^2^, 1111,8 mW/ cm^2^, 1209,9 mW/ cm^2^.) for 3 min. The entire experimentation was carried out in the dark box to avoid the influence of the ambient light. After irradiation, the experimental samples were incubated at 37°C for 48 hours. And then the growth situation of bacteria in different groups was observed and recorded. The detailed experiment setup is shown in Figure 1.

**Fig 1.**
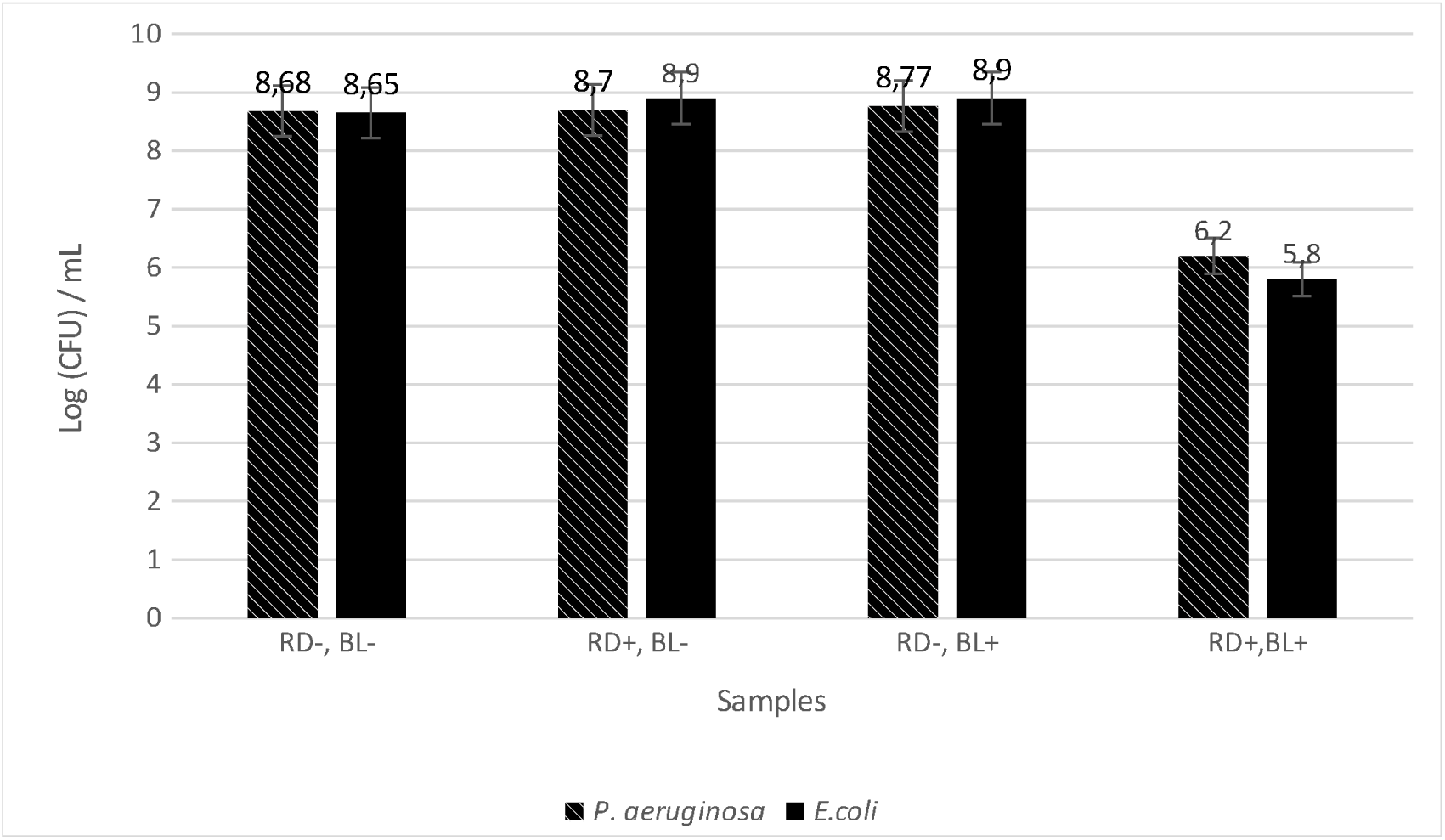
The number of survival bacterial colonies under different conditions of experiment (R: riboflavin (I); BL: blue light) to *E. coli* and (light power density: 1024,2 mW/ cm^2^)

### Bacteria Inactivation Experiments In Vitro (1) Inhibition Effect of Different BL Energy

0.5 MCF bacteria suspensions of *E. coli* and *P. aeruginosa* diluted into 1:10 with PBS were taken as BL alone group samples. 100 µL bacteria suspensions (0.5 MCF) and 400 µL RD solution (2.5% wt) were taken into 500 µL PBS as BL-RD group samples. The initial population densities of experimental samples were both maintained at about 10^9^ CFU/ml and the concentration of RD solution was about 0.02%wt. 150 µL aliquots of the sample solutions were subjected to 96-well culture plates (BioRad, New York, USA). The inside diameter of the well is about 5 mm. The distance of light source to surface of solution was 5 mm. And the light spot diameter was about 7 mm at this distance, which was larger than the well and enabled satisfying irradiation of suspension surface with BL. The illumination was conducted in dark box to prevent photosensitization of riboflavin from background visible light. The energy density was 912,501 mW/ cm^2^, 1024,209 mW/ cm^2^, 1209,920 mW/ cm^2^, respectively, and the irradiation time was 3 min. For the control group samples, the suspensions in 96-well plates were kept in the dark at room temperature for 3 min.

### Cell Concentration Determination and Survival Fraction Calculation

After the experimental treatments, aliquots (100 µL) were withdrawn from each well and serially diluted 10-fold with PBS. Ten microliters from each dilution mixture was streaked onto blood agar plates (BioRad, New York, USA) in triplicate. After incubation for 48 hours at 37°C, bacterial colonies were counted.

The survival fraction was calculated according to the equation *N*/No., where No. is the number of CFU / mL of bacteria without being treated and *N* is the number of CFU / mL of bacteria treated with BL and RD. All results were presented as means ± standard deviation (SD) of at least three independent experiments and each was measured in triplicate.

### Statistical Analysis

Experimental data were confirmed to be of normal distribution by the *K*-*S* test. Descriptive statistics were used to summarize the data in multiplex analyses. The results were shown as the means ± SD. The statistical significance between groups was determined using two-way analysis of variance (ANOVA). *P* < 0.05 were considered to be statistically significant.

## Results & discussion

In the first experiment, it was necessary to select the optimal radiation power, which would effectively inactivate the bacterial suspension for 3 minutes of exposure time. With a longer exposure time, the solution begins to heat up significantly. Table 1 shows the dependence of the percentage of surviving bacteria based on the lamp radiation power.

As can be seen from table 1. the most effective was irradiation of the solution at a flow rate of 1024.2-1111.8 mW / cm2. For further experiments, we chose a power of 1024.2 mW / cm2, during which some single colonies still survive and this power represents the lowest limit of efficiency.

The second experiment was to demonstrate the ability of a new dynamic derivative with its minimum content in solution to inactivate the experimental bacterial suspensions under the influence of visible light. Fig. 1 shows an experiment with pure riboflavin as a control substance.

**Table 2.** Protection properties of photo inactivated and *E. coli* in mice model as pseudomonas and esherichiosis peritonitis (IP injection 1 mL 4 log (CFU)/mL microorganisms suspension per mouse)

| N | Group/vaccine dose, mL,<br>4 log (CFU)/mL | Mice survival under vaccination by RD-BL inactivated bacteria, n=20 |  |
| --- | --- | --- | --- |
|  |  | *P. aeruginosa* (A) | *E. coli* (B) |
| 1 | Control* | 4 | 6 |
| 2 | 0,05 | 4 | 7 |
| 3 | 0,1 | 9<sup>1</sup> | 15<sup>1</sup> |
| 4 | 0,5 | 20<sup>1,2,3</sup> | 20<sup>1,2,3</sup> |
| 5 | 1,0 | 20<sup>1,2,3</sup> | 20<sup>1,2,3</sup> |
|  |  | Mice survival under vaccination by formalin inactivated bacteria (FIB), n=20 |  |
| 6 | 0,05 | 3 | 4 |
| 7 | 0,1 | 3 | 5 |
| 8 | 0,5 | 7<sup>1,3</sup> | 9<sup>1,3</sup> |
| 9 | 1,0 | 12<sup>1,3</sup> | 14<sup>1,3</sup> |

\*- vaccination by 0,9% sol. NaCl
<sup>1</sup>- statistically differences from Control, (P<0,05)
<sup>2</sup>- statistically differences from corresponding groups (4 and 8, 5 and 9), (P<0,05)
<sup>3</sup>- statistically differences from corresponding groups (4,5 and 8, 9), (P<0,05)

As can be seen from Figure 1, control samples without BL irradiation and not containing riboflavin (R-, BL-) remained viable in the range of 8.7-8.6 Log (CFU) / mL. The samples containing riboflavin that were not irradiated with BL (R +, BL-) did not statistically differ from the control group. Samples not containing riboflavin but irradiated with BL (R-, BL +) showed a statistically insignificant upward trend in (8.77–8.9) Log (CFU) / mL. A statistically significant decrease in the number of live bacteria was observed only for the sample containing riboflavin and irradiated with BL and amounted to (6.2-5.8) Log (CFU) / mL, respectively. In Figure 2, we present a similar experiment with a dynamic succinylated riboflavin derivative with incompletely substituted hydroxyl groups with respect to the carbohydrate component RD. Due to the presence of residues of carboxyl groups in different positions and combinations, the molecule penetrates better into the cell, and the lability of the ester bond leads to the formation of pure riboflavin and free succinic and maleic acids inside the cell. This allows you to significantly enhance the effectiveness of photodynamic inactivation at the same concentration of the photodynamic agent RD.

As can be seen from Figure 2, control samples without BL exposure and not containing RD remained viable in the range of 8.7-8.6 Log (CFU) / mL. Samples containing RD but not irradiated with BL (RD +, BL-) did not statistically differ from the control group. A sample not containing RD but irradiated with BL (RD-, BL +) again showed a statistically insignificant tendency to increase (8.6-8.9) Log (CFU) / mL, but only for E. coli. A statistically significant decrease in the number of live bacteria almost to complete inactivation was observed only for a sample containing RD and irradiated BL and in the range of (0.32-0.12) Log (CFU) / mL, respectively.

0.02% RD was capable of inactivating 6 orders magnitude more of bacteria (6 log CFU), than R at the same concentration. Then a suspension of *P. aeruginosa* and *E. coli* 4.0 Log (CFU) / mL was prepared, RD was added to a final concentration of 0.02%, and BL solutions were irradiated for 5 minutes at a power of 1024.2 mW / cm^2^. When the solution was irradiated for 5 minutes, the growth of bacteria was not determined (not shown in the graph). Inactivated suspensions were further used as particulate vaccines to study their effectiveness versus similar vaccines obtained by formalin treatment. Table 2 presents the results of a study of the protective properties of these vaccines.

As can be seen from table 2, in the control group, instead of vaccines, 1.0 ml of Normal saline was given intravenously. The 2 weeks later after infection was administered to control group, only 4 mice survived from 20 animals in group A, and only 6 animals in group B. Vaccination of animals with vaccine in doses of 0.05 ml 4 log (CFU) / mL for the experimental study of RD-BL and for the variant with formalin inactivated bacteria (FIB) did not statistically differ from the control group. For formalin-inactivated bacteria vaccines, a dose of 0.1 ml of 4 log (CFU) / mL also did not lead to a statistically significant increase in mouse life expectancy. For the experimental group, inactivated RD-BL, the dose of 0.1 ml statistically significantly differed both from the control and from the same dose of FIB vaccines. An increase in vaccine doses up to 0.5 ml gave more significant statistical differences not only from the control group, but also between the RD-BL and FIB groups. If a dose of 0.5 ml of 4 log (CFU) / mL was used in the RD-BL group, all animals in group A and group B survived. For a dose of 1 ml of 4 log (CFU) / mL in the RD-BL experimental group, all animals also survived. Whereas for formalin inactivated bacteria (FIB) groups, a 100% protective effect was not achieved, although the number of surviving animals was 4 times greater than in the control.

**Fig 2.**
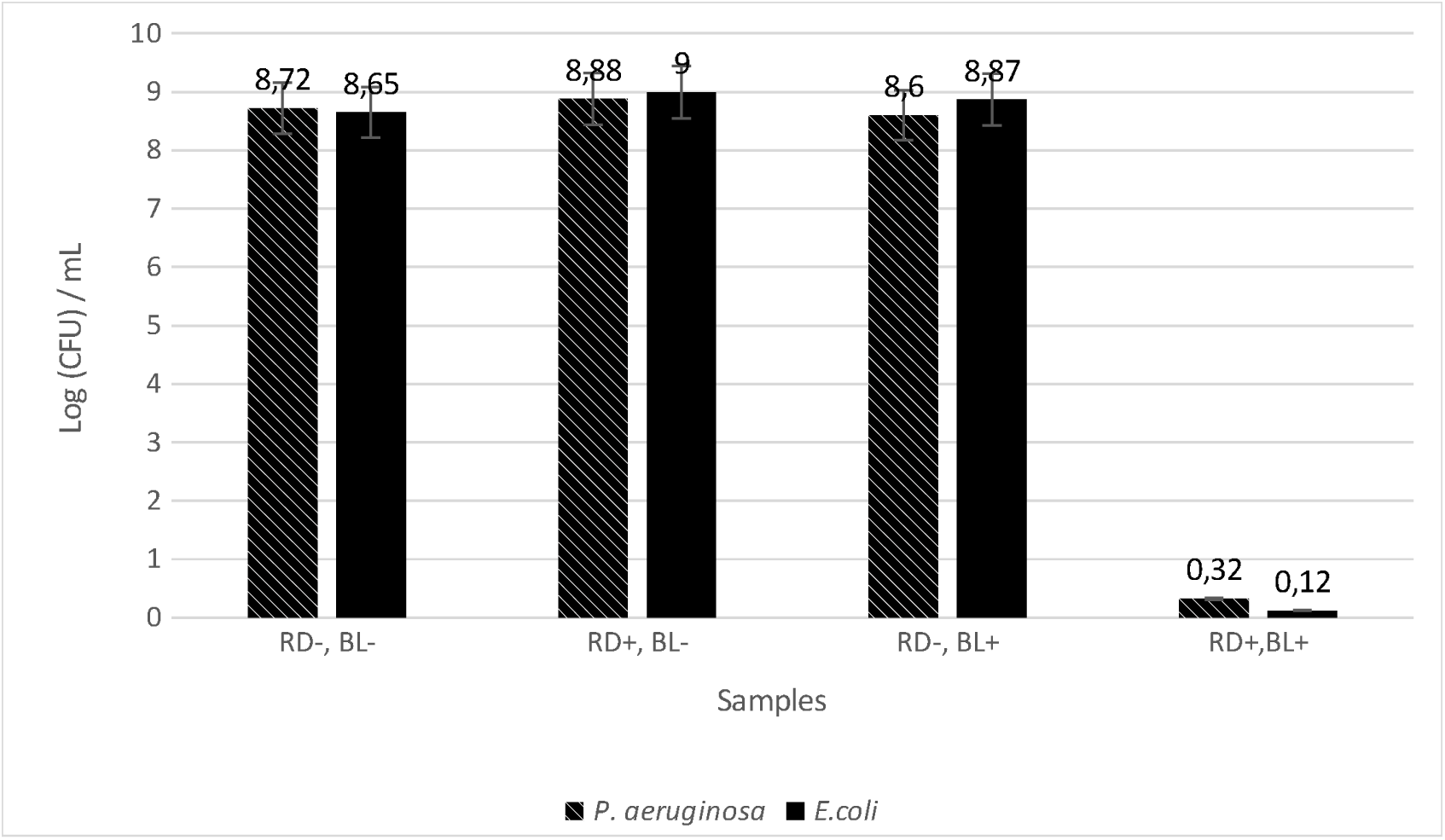
The number of survival bacterial colonies under different conditions of experiment (RD: riboflavin derivative; BL: blue light) to *E. coli* and (light power density: 1024,2 mW/ cm^2^)

## Conclusions

1. The dynamic derivative of riboflavin RD based on combinatorial succinylated - maleated riboflavin has photodynamic microbicidal activity in the visible irradiation spectrum with respect to *P*.*aeruginosa* and *E*.*coli*
2. 0.02% RD can photoinactivate 6 time more bacteria P.aeruginosa and E. coli than riboflavin.
3. The minimum effective BL power (450 nm) for the photodynamic inactivation of RD of both *P*.*aeruginosa* and *E. coli* is 1024.2 mW / cm2, irradiation of which for 3 minutes leads to the survival of some single colonies.
4. A 5-minute irradiation of bacterial suspensions of P. aeruginosa and E. coli BL in the presence of RD at a power of 1024.2 mW / cm2 leads to a complete loss of bacterial viability.
5. In groups of mice pre-vaccinated intraperitoneally with corpuscular vaccines BL-RD based on suspensions of *P*.*aeruginosa* and *E. coli* at doses of 0.5-1.0 ml 4 log (CFU) / mL, 100% survival of all animals was observed, whereas in control group with the same type of vaccines but formalin-treated vaccines, a 100% protective effect was not achieved.

